# Comparative methods for iPSC-Derived endothelial cells in modeling vascular diseases

**DOI:** 10.64898/2026.08.20.746033

**Authors:** Pakize Nur Akkaya, Louet Koolen, Zohreh Hosseinzadeh

## Abstract

Endothelial cells (ECs) derived from human induced pluripotent stem cells (hiPSCs) are increasingly used to model vascular diseases and test therapeutic strategies. However, the efficiency and reproducibility of differentiation can vary depending on the culture medium and its supplemented factors and stages. Here, we directly compared two defined media, APEL and BPEL, for iPSC-to-ECs differentiation. iPSCs were differentiated over 10 days with sequential growth factor induction, followed by magnetic-activated cell sorting or flow cytometry for CD31^+^ cells. Both media produced ECs with similar morphology and marker expression, including CD31 and VE-cadherin. Functional assays demonstrated comparable tube formation, indicating equivalent endothelial functionality. Cost analysis indicated that APEL had a higher total reagent cost but generated a higher total cell yield, resulting in a comparable cost per 10⁶ total cells, whereas BPEL was more cost-efficient for producing CD31⁺/VE-cadherin⁺ endothelial-specific cells. Our results suggest that APEL and BPEL media are equally effective for generating iPSC-derived ECs, providing flexibility in method selection for vascular disease modeling and drug discovery applications.

**Graphical abstract:** 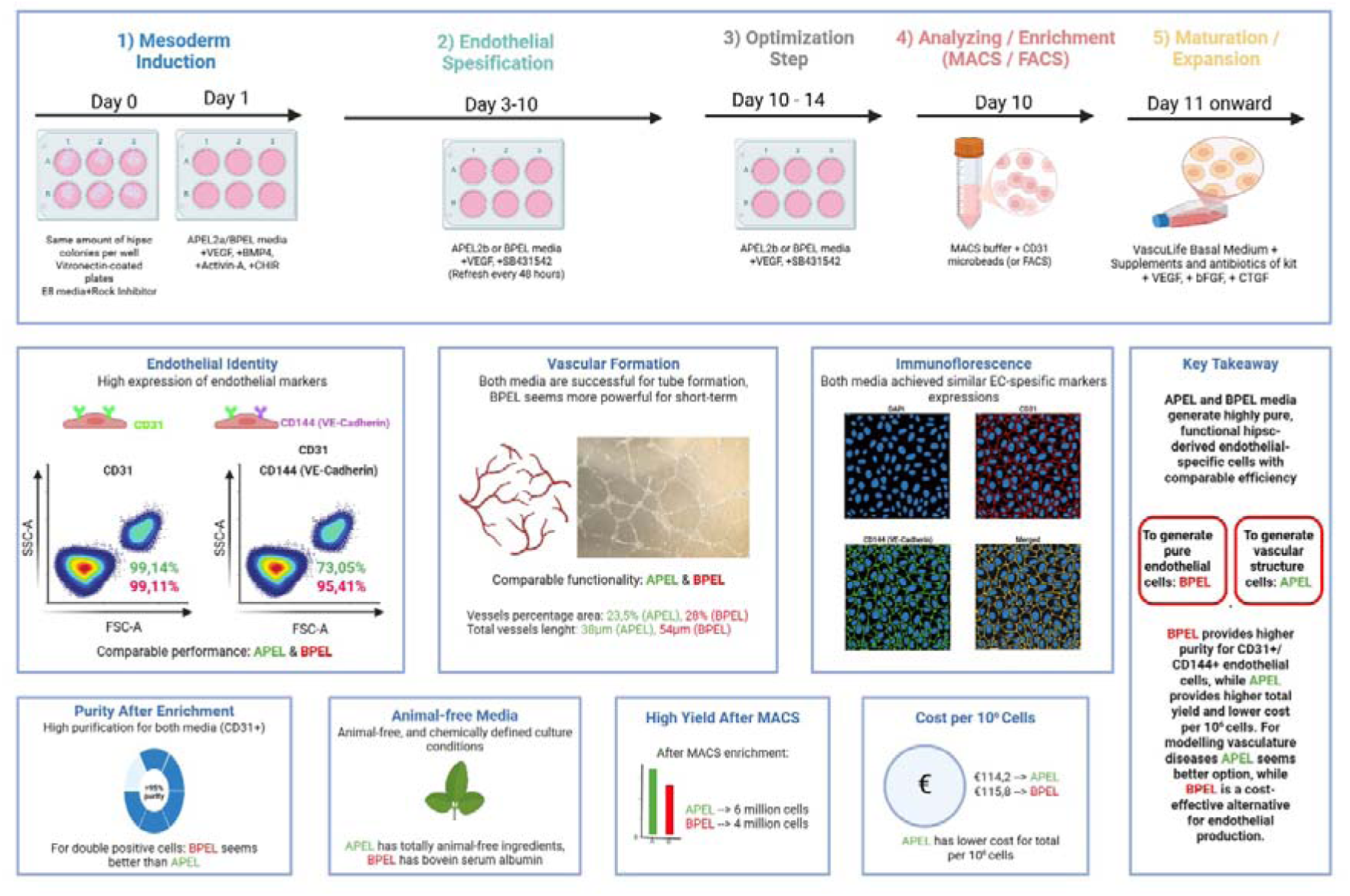

## Background

### Stem cells-derived ECs

Induced-pluripotent stem cells (iPSCs) and human induced-pluripotent stem cells (hiPSC) are pluripotent-ability cells via reprogramming. They are derived from somatic cells and possess to ability to differentiate into every cell types in the body [1]. In tissue engineering applications, specific cells derived from iPSCs are used with some biomaterials such as hydrogels and biomaterial composites like gelatin, collagen, hydroxiapatite etc. to obtain functional tissues [2]. For disease modelling, one of the most effective and widely used ways; especially for genetic diseases, is modelling patient-derived iPSCs, and this is quite an important platform to model the diesases, understanding the disease mechanisms, and drug target research [3].

Endothelial cells (ECs) are lining the inner surface of the blood vessels. They are important for vascular biology and responsible for vascular permeability, inflammation responses and angiogenesis [4]. In regenerative medicine area, these cells have a critical role for vascularization especially in tissue engineering to ensure vascularisation for tissue survival. [5]. In addition, iPSC-derived ECs are also used for treatment of ischemic tissues [6].

EC differentiation from stem cells can be achieved through three main strategies: (a) induction via 3D embryoid body (EB) formation, (b) directed differentiation in 2D monolayer cultures using specific signaling molecules, and (c) genetic programming through the introduction of key endothelial genes [7]. In the EB approach, hiPSCs are aggregated into 3D structures, followed by spontaneous or semi-conductive differentiation [8]. However, the ECs get from these protocols exhibit high heterogenity including arterial, veneous, lymphatic ECs. To address this, some studies employ more defined differentiation strategies targeting specific signaling pathways, such as BMP4 and VEGF-A, to guide lineage specification [9].

Around 2014, iPSC-derived endothelial differentiation protocols began to shift from three-dimensional embryoid body (EB)-based systems toward two-dimensional monolayer approaches. In these protocols, hiPSCs are first directed toward mesodermal lineage commitment and subsequently specified toward the vascular endothelial lineage. This staged 2D strategy improved protocol control, endothelial yield, reproducibility and scalability compared with earlier EB-based methods. [7]. Orlova et al. developed a directed differentiation protocol in which hiPSCs are first treated with BMP4, Activin A, and VEGF to induce mesoderm formation and initiate vascular specification [10]. This is followed by VEGF combined with a TGF-β inhibitor (SB431542), which promotes endothelial differentiation while suppressing alternative lineages [11]. Similarly, in another protocol using small molecules, the differentiation of hiPSCs through the mesodermal cells first, and then ECs was achieved with VEGF+forskolin [12]. The advantage of this protocol is that ECs can be obtained with high efficiency and purity in a short time. This procedure also has the advantage of generating vascular smooth muscle cells using with PDGF-BB, alongside ECs.

Another protocol using the combination of VEGF and cGMP has stimulated the production of hiPSC-derived ECs in 2D monolayer and serum-free conditions, while also eliminating cells that do not show early-stage endothelial differentiation. Thus, productivity reached nearly 100%, and this production was achieved without the need for EC sorting procedure [13]. The protocol that produces a defined 8-day protocol by activating WNT, initially promoting mesodermal differentiation, then endothelial differentiation with VEGF/FGF, and finally achieving pure hiPSC-derived EC isolation through CD144+ MAC sorting, is also significant in this historical timeline [14].

Recently, Xeno-free matrices, well-defined media, and 3D suspension/bioreactor variants have been developed. These protocols facilitated the production of better-defined, stable, and targeted hiPSC-derived ECsthat are more independent in terms of animal components and can be adapted for translational use [15]. A serum-free developed protocol, applicable in both 2D monolayer and 3D bioreactor suspension cultures, lasting a total of 6 days, enabled the production of hiPSC-derived ECs with approximately 90% efficiency and strong arterial characteristics. Consequently, hiPSC-derived ECs have commenced utilization for production in a translational context, moving beyond the only research-oriented domain [16].

Shorter differentiation protocols have recently been developed based on the overexpression of ETV2, which achieves approximately 99% purity within a very short period of 5 days without requiring cell sorting [17]. Similarly, alternative protocols that directly program hiPSC-derived endothelial cell progenitors using the combination of SOX17+FGF2 factors are also a good example of protocols developed with transcription factors in recent times [18].

Some new protocols that allows the production of both hiPSC-derived ECs, pericytes, and fibroblasts with a single differentiation protocol has been particularly valuable for laboratories working on vascularized organoids and multicellular model systems, as it offers a system that can also create a vascular microenvironment [19]. Today several research groups, including ours are focusing on producing organ-specific hiPSC-derived EC subtypes rather than general endothelium production. A strategy that specifically entails the generation of venous ECs enriched with retinoic acid and attains success through cell cycle modulation is remarkable in this area [20].

### Medium for iPSC-derived EC differentiation

In iPSC-derived EC differentiation, medium choice critically affects production and protocol optimization. Media can be classified into three categories: (a) basal media for mesoderm induction, (b) EC-specific media that promotes the mesoderm-to-endothelial transition, and (c) mature EC media that supports maintenance and expansion [6].

RPMI 1640 supplemented with B27 represents a simpler and more cost-effective alternative than standard basal media that includes basically albumin, polyvinylalcohol, and essential lipids and has become the most commonly used medium in WNT signaling-based differentiation protocols, largely owing to its widespread adoption in cardiac differentiation workflows. In this formulation, RPMI 1640 serves as the basal medium, whereas the serum-free B27 supplement provides insulin, antioxidants, vitamins, lipids, and other nutrients required for cell growth and differentiation [11]. Although RPMI/B27 effectively supports mesoderm induction, it was not specifically developed for endothelial differentiation. Despite its low cost and widespread use, RPMI/B27 was not specifically designed for endothelial differentiation and therefore generally requires additional optimization to achieve efficient EC specification. Among the endothelial-specific growth media used to expand and maintain the EC population obtained after mesodermal induction and endothelial differentiation stages, the most common is the EGM-2 (endothelial growth medium-2) kit. This medium kit contains endothelial basal medium-2 and added growth factors and supplements such as VEGF, bFGF, IGF, EGF, Hydrocortisone, ascorbic acid, heparin, and fetal bovine serum (FBS) [19]. Serum-free, chemically defined media are generally preferred for iPSC-derived EC differentiation because they reduce batch-to-batch variability, enable precise modulation of developmental signaling pathways, and improve the reproducibility and clinical translatability of differentiation protocols. In this context, serum-free EC maintaining media have also started to be used in recent years. Endothelial SFM (serum-free medium), xeno-free custom media, and some STEMdiff products are examples of such media. these media supplements with several factors for EC differentation including, VEGF, FGF2, IGF1, recombinant albumin, transferrin, and defined lipid contents have been added to a basal endothelial medium [15].

The selection of medium for the production of iPSC-derived ECs is extremely important in terms of both yield and purity as well as cost effectivness. Although well-defined, standardized, and serum-free media such as APEL are recognized for their superior efficiency, purity, and repeatability, their frequently elevated cost can impose financial burdens on laboratories, particularly in long-term, repeated, and sustainable research endeavors [10]. Simultaneously, StemPro-34 is among the more costly media used for iPSC-derived EC differentiation. Although it can initially support high EC yields, prolonged use or application outside the optimal differentiation window may compromise endothelial purity. Thus, medium selection should be guided by the stage-specific requirements of the differentiation protocol, rather than by the assumption that higher-cost media necessarily produces superior outcomes [21]. The cost of differentiation media is not a secondary consideration but a recognized barrier to achieve a scalable hiPSC-derived EC production, highlighting the need for efficient and cost-effective protocols to enable large-scale clinical translation [14, 16].

APEL (Albumin, Polyvinylalcohol, Essential Lipids) medium and BPEL (Bovine Serum Albumin, Polyvinylalcohol, Essential Lipids) medium are commonly used basal media for initiating iPSC-derived EC differentiation. APEL is a commercially available, chemically defined, serum-free and animal-origin-free medium that provides high reproducibility and is compatible with both embryoid body-based and monolayer differentiation systems, making it suitable for translational applications [22]. Although its exact formulation is proprietary, APEL contains recombinant albumin, insulin, transferrin, lipid supplements, essential amino acids and vitamins in a DMEM/F12-like basal formulation [22]. By contrast, BPEL can be prepared in-house and is therefore substantially more cost-effective than APEL, but it contains bovine serum albumin (BSA) and polyvinyl alcohol (PVA). Both media are typically supplemented with morphogens and growth factors, including BMP4, VEGF, CHIR99021 and bFGF, to promote mesoderm induction and endothelial specification [10]. Because BPEL contains animal-derived components, it may be less suitable than APEL for clinical or xeno-free applications. In addition, its BSA content may introduce greater batch-to-batch variability and reduced formulation consistency compared with recombinant albumin-based media [23–25]. These limitations may explain why BPEL is less frequently selected for long-term or translational studies despite its economic advantages.

Despite the widespread use of APEL and BPEL in iPSC-derived EC differentiation protocols, the extent to which their compositional differences influence endothelial differentiation efficiency remains unclear, leaving an important gap in identifying the most appropriate basal medium for research-scale EC differentiation from iPSCs. In this study, we directly compared APEL and BPEL for EC differentiation. Our results show that BPEL is a cost-effective alternative to APEL that supports comparable EC differentiation and preserves key EC characteristics. Notably, flow cytometric analysis revealed a higher proportion of CD31⁺/VE-cadherin⁺ hiPSC-ECs in BPEL-derived cultures than in APEL-derived cultures.

## Material and Method

### Protocol for EC differentiation from iPSCs with APEL medium

#### 1. iPSC Culture and Seeding

1. Maintain undifferentiated iPSC until ∼80% confluency (split ratio 1:3 or 1:4).
2. On the day cells reach the desired confluency (Day 0):
  - Harvest cells with 1mL/per well 0.5M EDTA solution, following 37°C / 5 min incubation, tap the wells, discard EDTA, and resuspend the colonies in E8 medium and plate onto Vitronectin-coated 6-well plates.
  - Coat each well with 1 mL Vitronectin solution (10 µL of 0.5 mg/mL Vitronectin in 1 mL PBS) and incubate it for 1 hour at Room Temperature (RT)
  - Add 2 mL E8 medium supplemented with 5 µM Rock inhibitor per well.
  - Plate across 12–18 wells as needed.

#### 2. Differentiation induction

**Day 1:**

1. Remove E8 medium.
2. Add 2.5 mL APEL2a medium per well (APEL2a: APEL2 medium + 50 ng/mL VEGF165, 30 ng/mL BMP4, 25 ng/mL Activin-A, 1.5 µM CHIR). (Fig. 1)

**Fig. 1:**
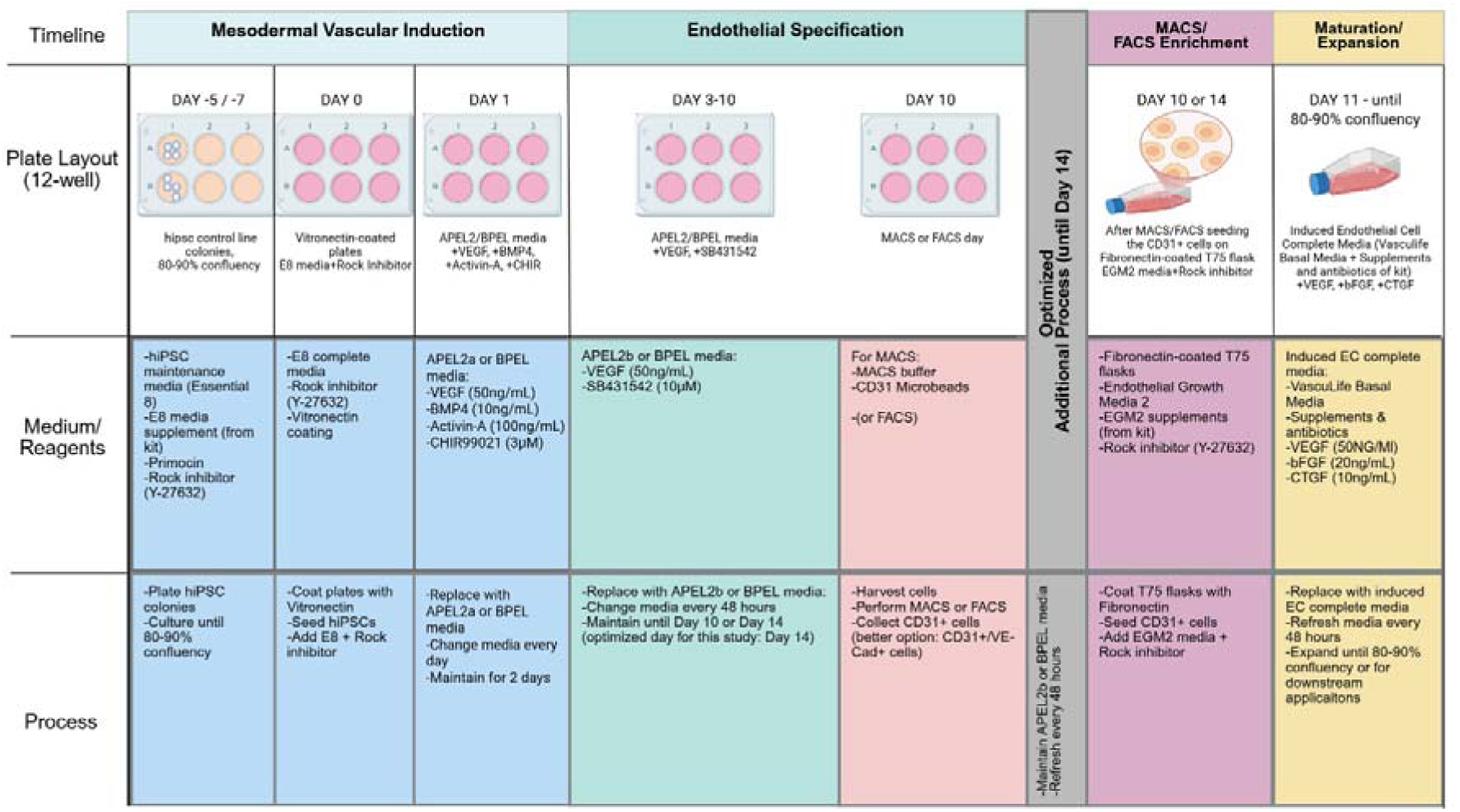
Schematic process of hiPSC-derived ECs generating with APEL or BPEL media. The protocol begins with maintenance of hiPSC colonies until they reach around 80–90% confluency. Cells are then plated on vitronectin-coated wells in E8 medium with ROCK inhibitor. On day 1, mesodermal and vascular induction is initiated using APEL2 or BPEL medium supplemented with VEGF, BMP4, Activin-A, and CHIR99021. From day 3 to day 10, endothelial specification is promoted using APEL2 or BPEL medium containing VEGF and SB431542. On day 10 or day 14, differentiated cells are enriched for endothelial cells by MACS or FACS, mainly targeting CD31-positive cells. After enrichment, cells are seeded onto fibronectin-coated flasks in endothelial growth medium. Finally, the purified endothelial cells are matured and expanded in induced endothelial cell complete medium until they reach 80–90% confluency for downstream experiments.

**Day 3:**

1. Remove APEL2a medium.
2. Add 2.5 mL APEL2b medium per well (APEL2b: APEL2 medium + 50 ng/mL VEGF165 + 10 µM SB431542).

**Day 5:**

Refresh APEL2b medium (2.5 mL per well).

**Day 7–10:**

1. Either perform MACS sorting on Day 7 or extend differentiation to Day 10 (preferred based on experience).
2. If extending, refresh APEL2b medium (3 mL per well) (Fig.2).

**Fig. 2:**
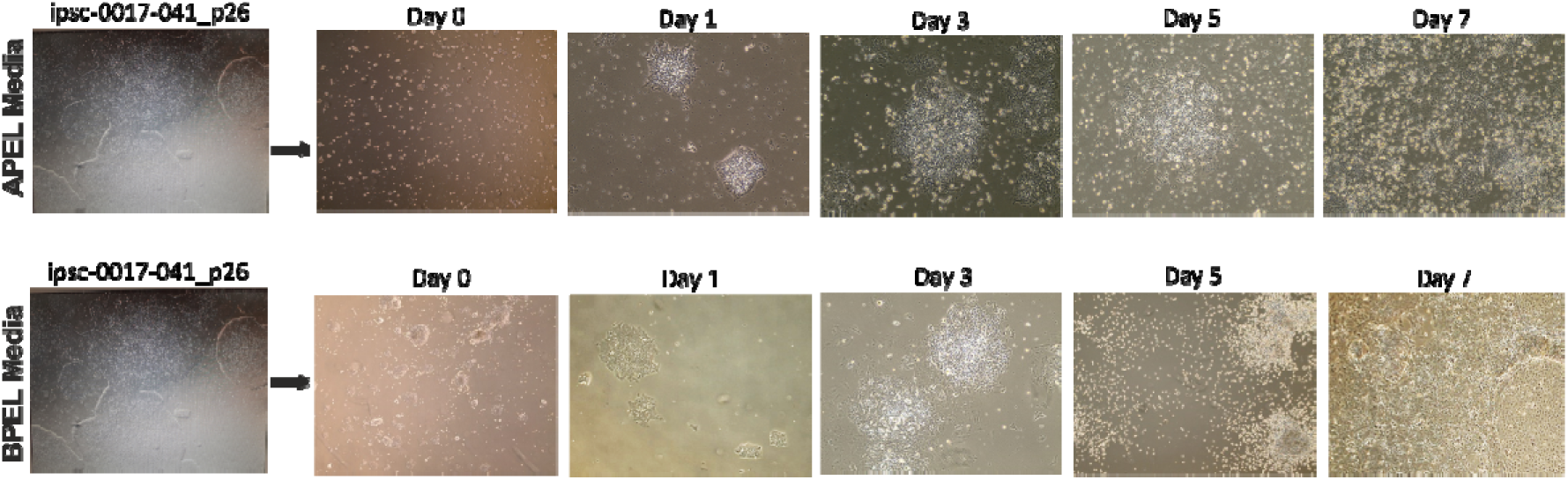
hiPSC-derived EC Differentiation using APEL and BPEL media. Representative phase-contrast images showing the morphological changes during differentiation of human iPSCs into EC. Undifferentiated iPSC colonies at high confluency prior to induction (Day 0), followed by early differentiation stages after mesoderm induction (Day 1–3), characterized by cell dispersion and loss of compact colony structure, and the emergence of loosely organized cell clusters,progressive EC differentiation (Day 5–7), showing the formation of dense cell aggregates and increased cell proliferation, followed by the appearance of more uniform, cobblestone-like morphology typical of ECs at later stages in APEL (first row) and BPEL media (second row).

#### 3. Magnetic-activated cell sorting (MACS) for CD31+ cells (Day 10)

1. Wash cells with 2 mL PBS per well.
2. Incubate with 0.5 mL TrypLE per well at 37°C for 5 min.
3. Gently pipette to detach cells; incubate for an additional 5 min.
4. Collect cells into 10 mL cold MACS buffer and bring total volume to 45 mL with MACS buffer.
5. Filter cells through a 20–30 µm strainer.
6. Centrifuge at 1000 rpm for 5 min and remove supernatant.
7. Resuspend cells in 5 mL MACS buffer + 5 µM Rock inhibitor per 6-well plate.
8. Count cells (expected yield: ∼1.5 × 10^6 per well; e.g., 12 wells ≈ 18 × 10^6 cells).
9. Centrifuge again, remove supernatant, and resuspend in Dynabeads CD31 solution (35 µL MACS buffer + 25 µL Dynabeads per 10–100 million cells).
10. Incubate at 4°C in the dark for 15 min.
11. Wash with 20 mL MACS buffer, filter, and centrifuge (5 min, 1000 rpm).
12. Resuspend cells in 1 mL MACS buffer + 5 µM Rock inhibitor and prepare MACS column:
  - Wash column.
  - Load cells.
  - Collect CD31− fraction. Repeat washing with 3 mL MACS buffer if needed.
  - Remove column from stand.
  - Elute CD31+ cells with 2 mL EGM2 medium + 20 µL Revitacell.

#### 4. Post-MACS

1. Plate CD31+ cells at maximum 7.5 × 10^4 cells per fibronectin-coated T75 flask.
2. Next day, replace medium with induced Endothelial Cell (iEC) complete medium:
3. VascuLife Medium kit (basal medium + supplements + antibiotics)
  - VEGF165 (25 ng/mL)
  - bFGF (20 ng/mL)
  - CTGF (10 ng/mL)
4. Refresh medium every 2–3 days.
5. Passage cells at 80–90% confluency: wash with PBS, incubate with 5 mL TrypLE (5 min, 37°C), centrifuge, and resuspend in 1 mL iEC medium. Plate 5 × 10^5 cells per fibronectin-coated T75 flask or cryopreserve.

#### 5. Flow cytometry analysis after MACS

After MACS and seperating all cells as CD31(+) and CD31(-) cells into different two tubes in EGM2 + Revitacell media for CD31(+) cells, E8 flex + Revitacell media for CD31(-) cells:

1. Count them via automated cell counter.
2. Seperate 4-5 × 10^5^ cells from CD31(+) cells tube, and seperate 2-2,5 × 10^5^ cells from CD31(-) cells tube.
3. Wash these seperated cells with PBS (2 times).
4. Centrifuge them at 1000 rpm, 5 mins.
5. Discard supernatant.
6. Resuspend the cells in 100µL Flow Cytometry buffer per tubes.
7. Filter them with 70 µm filter, and put them into Flow Cytometry tubes.
8. Add 1/100 antibody (CD31-APC) to the tubes that you marked as CD31(+) and CD31(-). (Should be two more tubes for unstained control per each group, no antibody).
9. Incubate the tubes at 4°C in the dark, 20-30 mins.
10. Wash the cells with PBS.
11. Centrifuge them at 1000 rpm, 5 mins.
12. Resuspend them in 300-500 µL FC buffer per tubes.
13. Bring the tubes to the FC analyzing process.

#### 6. Flourescence-activated cell sorting (FACS)

After differentiation protocol, on Day 10 (or 14), FACS was applied without MACS for sorting CD31+ and CD31-cells, and also identifying the ratios of CD31+, CD31+/CD144+, CD31-cells. For this aim:

1. Wash the plates with PBS.
2. Harvest the all differentiated cells with TrypLE Express Enzyme (1X).
3. Do it for APEL-differentiated cells, and BPEL-differentiated cells seperately.
4. Collect the cells into APEL medium, and BPEL medium seperately.
5. Pipetting the cells gently.
6. Count the cells with viability dye via automated cell counter (Luna).
7. Centrifuge the cells at 1000 rpm, 5 mins
8. Discard supernatant per both tubes.
9. Resuspend the cells 100µL Flow Cytometry Buffer seperately.
10. Stain the cells with CD31-APC and CD31-APC/VE-Cad-488 dyes (1:100)
11. Incubate the tubes at 4°C, in the dark, 20-30 mins.
12. Wash the cells with FC buffer.
13. Centrifuge them at 1000rpm, 5 mins.
14. Discard supernatant per both tubes.
15. Resuspend them at 300-500µL FC buffer again.
16. Filter the cells with 70µm cell strainer to eliminate aggregates.
17. Put them into sorter tubes.
18. Bring the tubes to the FACS sorting and analyzing process.

#### 7. Buffers and media preparation

**MACS buffer** (store at 4°C, keep cold, work on ice):

- 25 mL BSA solution (5%)
- 1 mL EDTA (0.5 M)
- 474 mL PBS
- Filter sterilize

**EGM2 medium:**

- EGM2 Basal medium
- FBS 0.02 mL/mL
- rhEGF 5 ng/mL
- bFGF 10 ng/mL
- rhIGF1 20 ng/mL
- rhVEGF165 0.5 ng/mL
- Ascorbic acid 1 µg/mL
- Heparin 22.5 µg/mL
- Hydrocortisone 0.2 µg/mL

**Flow cytometry (FC) buffer:**

- 2% BSA
- PBS

### Protocol for EC differentiation from iPSCs with BPEL medium

#### 1. iPSC culture and seeding

1. Maintain undifferentiated iPSCs until ∼80% confluency (split ratio 1:3 or 1:4).
2. On the day cells reach the desired confluency (Day 0):
  - Harvest cells with 1mL/per well 0.5M EDTA solution, following 37°C 5 min incubation, tap the wells, discard EDTA, and resuspend the colonies in E8 medium and plate onto Vitronectin-coated 6-well plates.
  - Coat each well with 1 mL Vitronectin solution (10 µL of 0.5 mg/mL Vitronectin in 1 mL PBS) and incubate it for 1 hour at RT.
  - Add 2 mL E8 medium supplemented with 5 µM Rock inhibitor per well.
  - Plate across 12–18 wells as needed.

#### 2. Differentiation induction

**Day 1:**

1. Remove E8 medium.
2. Add 2.5 mL BPEL medium per well (BPEL medium supplemented with 50 ng/mL VEGF165, 30 ng/mL BMP4, 25 ng/mL Activin-A, 1.5 µM CHIR). (Fig. 1)

**Day 3:**

1. Remove BPEL medium.
2. Add 2.5 mL fresh BPEL medium per well (BPEL supplemented with 50 ng/mL VEGF165, 10 µM SB431542).

**Day 5:**

Refresh BPEL medium (2.5 mL per well).

**Day 7–10:**

1. Perform MACS on Day 7 (as recommended in reference protocols) or extend differentiation to Day 10 (based on experience).
2. If extending, refresh BPEL medium (3 mL per well) (Fig.3).

**Fig. 3:**
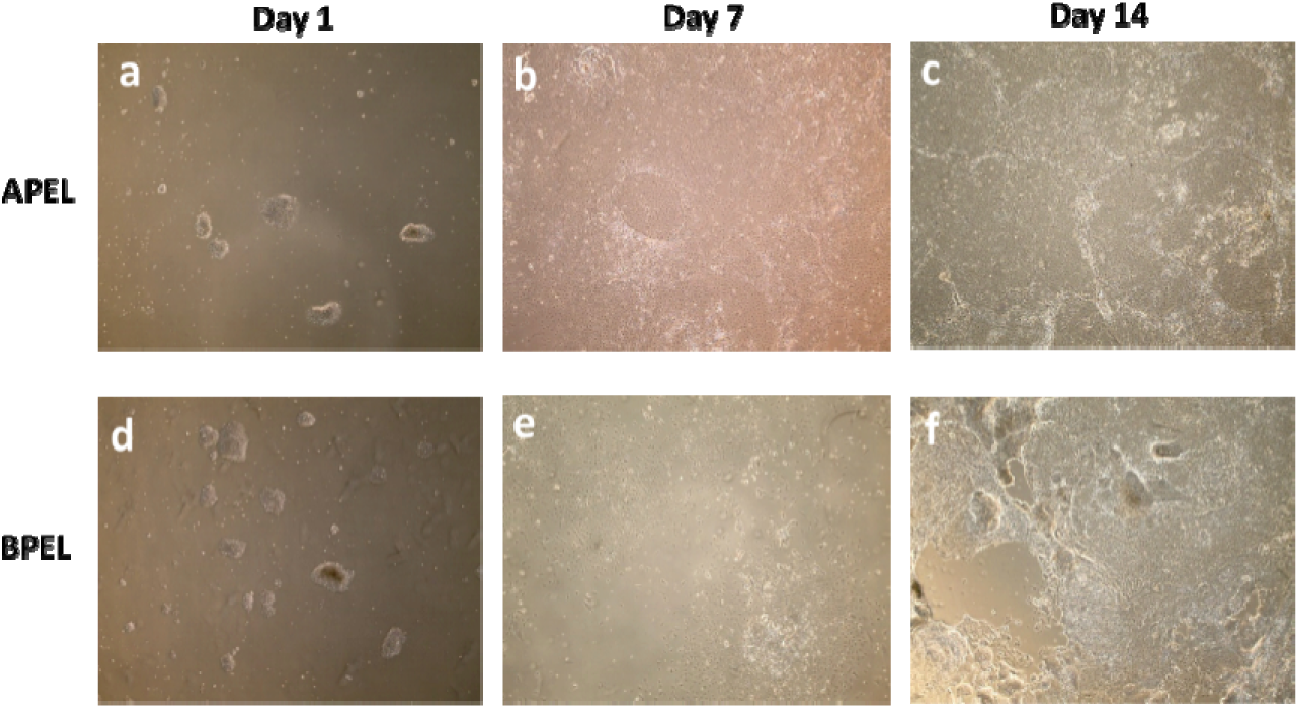
Comparison of APEL and BPEL media for EC differentiation. Representative phase-contrast images showing the morphological changes during differentiation of human iPSCs into ECs. Cells were imaged at day 1, 7, and 14 post-differentiation in APEL upper row (a,b, and c), and BPEL (lower row) (d, e, and f).

#### 3. MACS for CD31+ Cells (Day 10)

1. Wash cells with 2 mL PBS per well.
2. Incubate with 0.5 mL TrypLE per well at 37°C for 5 min.
3. Pipette gently, incubate for an additional 5 min.
4. Collect cells into 10 mL cold MACS buffer; bring total volume to 45 mL with MACS buffer.
5. Filter through a 20–30 µm strainer.
6. Centrifuge at 1000 rpm for 5 min; remove supernatant.
7. Resuspend cells in 5 mL MACS buffer + 5 µM Rock inhibitor per 6-well plate.
8. Count cells (expected yield: ∼1.5 × 10^6 per well; e.g., 12 wells ≈ 18 × 10^6 cells).
9. Centrifuge again, remove supernatant, and resuspend in Dynabeads CD31 solution (35 µL MACS buffer + 25 µL Dynabeads per 10–100 million cells).
10. Incubate at 4°C in the dark for 15 min.
11. Wash with 20 mL MACS buffer, filter, centrifuge, and resuspend in 1 mL MACS buffer + 5 µM Rock inhibitor.
12. Prepare MACS column:
  - Wash the column.
  - Load cells; collect CD31− fraction. Repeat washing if necessary.
  - Remove column from stand.
  - Elute CD31+ cells with 2 mL EGM2 medium + 20 µL Revitacell.

#### 4. Post-MACS

1. Plate CD31+ cells at a maximum of 7.5 × 10^4 cells per fibronectin-coated T75 flask.
2. Next day, replace medium with induced Endothelial Cell (iEC) complete medium:
  - VascuLife Medium kit (basal medium + supplements + antibiotics)
  - VEGF165 (25 ng/mL)
  - bFGF (20 ng/mL)
  - CTGF (10 ng/mL)
3. Refresh medium every 2–3 days.
4. Passage cells at 80–90% confluency: wash with PBS, incubate with 5 mL TrypLE (5 min, 37°C), centrifuge, resuspend in 1 mL iEC medium. Plate 5 × 10^5 cells per fibronectin-coated T75 flask or cryopreserve.

#### 5. BPEL medium composition

- IMDM (no phenol red)
- Ham’s F12 + Glutamax
- Protein-free hybridoma medium
- Chemically defined lipids
- ITS-X
- Glutamax
- Primocin
- BSA solution
- PVA solution
- 1-Thioglycerol (a-MTG)
- Ascorbic acid

## Troubleshooting

Several technical challenges may arise during the differentiation of iPSCs into endothelial cells using APEL and BPEL protocols. A common issue is low differentiation efficiency, often resulting from poor iPSC quality, over-confluency prior to induction, or reduced activity of key growth factors such as VEGF, BMP4, and Activin A. To minimize this, it is critical to use healthy, undifferentiated iPSCs at 70– 80% confluency and to prepare fresh aliquots of growth factors while avoiding repeated freeze–thaw cycles. Precise timing of media changes is also essential, as deviations can disrupt mesoderm induction and endothelial specification.

Cell death during early differentiation may occur due to insufficient use of ROCK inhibitor or harsh dissociation procedures. This can be mitigated by including ROCK inhibitor during seeding and minimizing mechanical stress during cell handling. Poor cell attachment is often linked to inefficient coating of culture surfaces; therefore, freshly prepared vitronectin or fibronectin coatings and appropriate incubation conditions are recommended.

During MACS-based CD31⁺ cell isolation, low yield may result from insufficient endothelial differentiation or suboptimal bead binding. Extending differentiation to Day 10 and optimizing Dynabeads concentration and incubation conditions can improve recovery. Additionally, excessive cell loss during washing or filtering steps should be avoided by gentle handling and appropriate centrifugation speeds. Contamination with CD31⁻ cells may indicate incomplete magnetic separation or column overloading, which can be addressed by repeating the sorting step or reducing cell input.

Functional deficiencies, such as poor tube formation or barrier integrity, may reflect incomplete endothelial maturation or suboptimal culture conditions. Prolonged culture in endothelial maintenance medium supplemented with VEGF and bFGF can enhance maturation. Finally, experimental variability between replicates or protocols (APEL vs BPEL) may arise from batch-to-batch differences in reagents, variability between iPSC lines, or sensitivity to signaling cues. Standardizing reagents, validating new batches, and optimizing growth factor concentrations for each condition are essential to ensure reproducibility and consistent outcomes.

## Results

### Comparison of APEL and BPEL media for iPSC-derived EC differentiation

The contrast microscopy for EC differentiation process from day 0 to day 7 did not show any major morphological differences between APEL and BPEL medium culture cells (Fig. 2 and 3). However in spite of starting with approximately the same amount of hiPSC colonies using both culture media, on MACS/FACS day (Day 10), there were approximately 6 million cells in APEL medium culture, but there were approximately 4 million cells in BPEL medium culture. According to the differentiated cells of flow cytometry analysis, the pre-sort population of CD31+ cells in APEL medium was 45,09%, and the pre-sort population of CD31+ cells in BPEL medium was 52,22% (Fig. 4). Flow cytometry analysis of endothelial cells differentiated in APEL medium and enrichedvia FACS demonstrated a highly pure population of CD31⁺ cells. Specifically, 99.14% of the sorted cells expressed the endothelial marker CD31. Within this CD31⁺ population, 73% co-expressed VE-Cadherin (CD144), confirming the endothelial identity and suggesting a substantial fraction of cells had acquired mature endothelial characteristics. These data indicate that APEL medium efficiently generates endothelial cells with robust marker expression following FACS enrichment (Fig. 5). Flow cytometry analysis of endothelial cells differentiated in BPEL medium and enriched via FACS demonstrated a highly pure CD31⁺ population, with 99.11% of cells expressing CD31. Among these CD31⁺ cells, 95% co-expressed VE-Cadherin (CD144; VE-Cad-488), confirming endothelial identity and indicating that the majority of cells had acquired mature endothelial characteristics. These findings indicate that APEL medium efficiently supports the generation and enrichment of functional endothelial cells comparable to BPEL medium (Fig. 5). On the other hand, according to the flow cytometry analysis after manually MACS protocol (there was no flow cytometry sorting step, there was just analyzing for manually seperated CD31+ cells), despite of expecting all of them are CD31+, just 74,95% of cells expressing CD31 (Fig. 6).

**Fig. 4:**
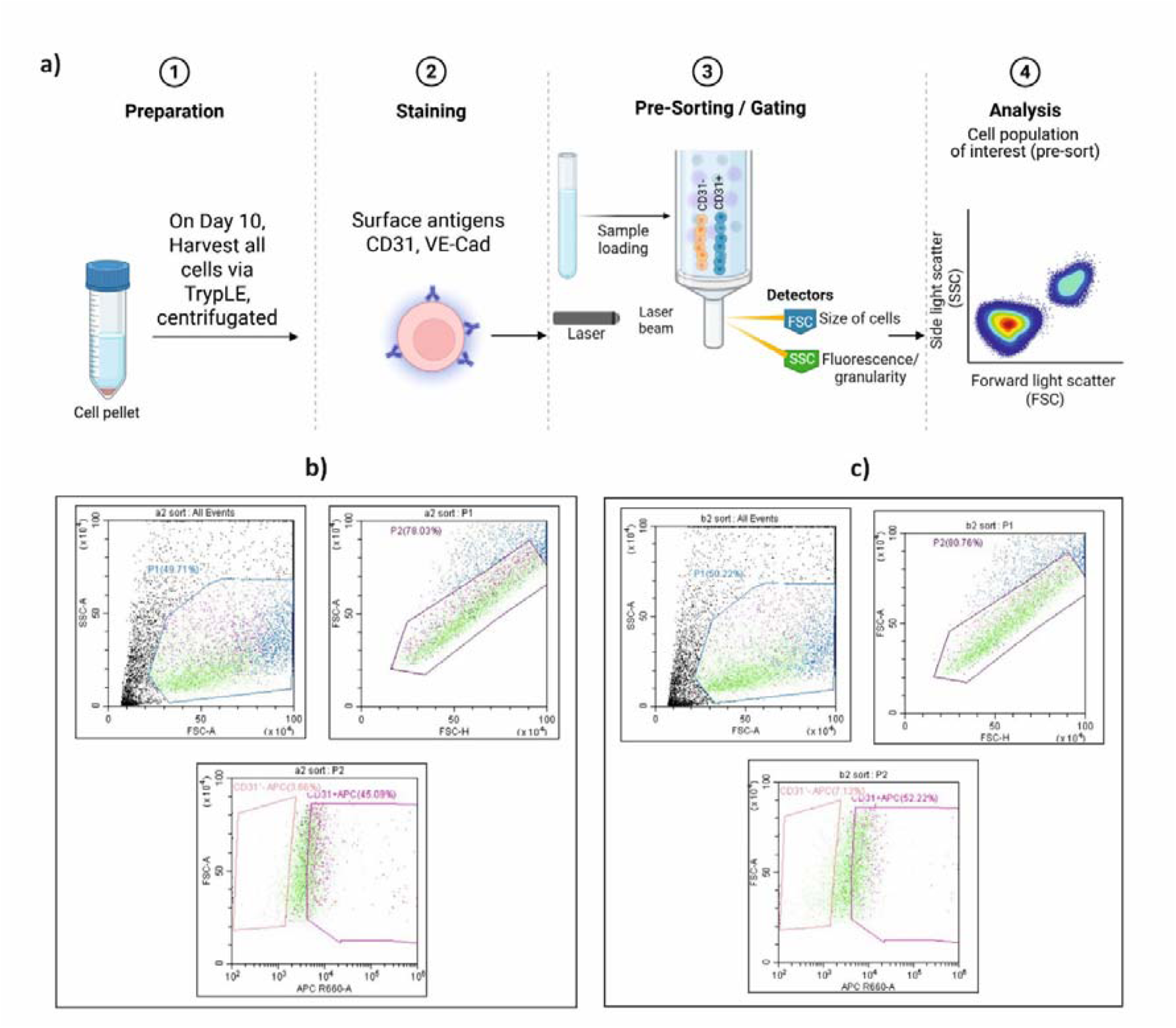
Flow cytometry analysis of EC markers in pre-sorting process for APEL and BPEL media. (a) Schematic figure of FACS sorting until gating process after EC differentiation at day 10. (b) EC were cultured in APEL media, were analyzed for expression of the EC marker CD31 pre-sorting (before fluorescence-activated cell sorting (FACS) enrichment with 45.09% of cells positive for CD31. (c) EC were cultured in BPEL media, cells were analyzed for EC marker CD31, in pre-sorting process, achieving with 52.22% CD31⁺ cells.

**Fig. 5:**
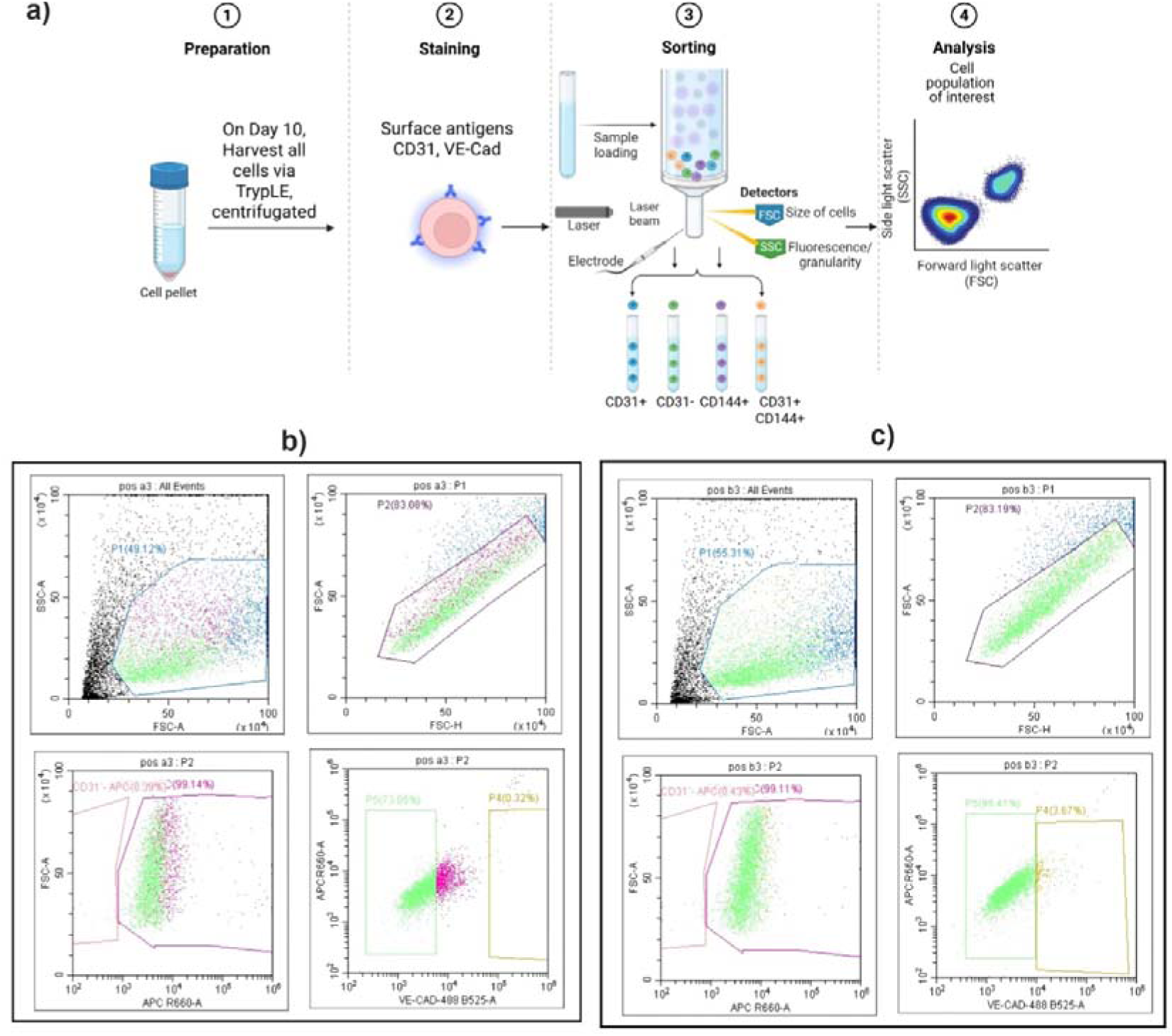
Flow cytometry analysis of EC markers following FACS enrichment for APEL and BPEL media. (a) Schematic figure of FACS sorting after EC differentiation at day 10. (b) EC were cultured in APEL media, were analyzed for expression of the EC marker CD31 after fluorescence-activated cell sorting (FACS). A high purity population was obtained, with 99.14% of cells positive for CD31. Among the CD31⁺ population, 73.05% co-expressed VE-Cadherin (CD144; VE-Cad-488). (c) EC were cultured in BPEL media, Cells were sorted by FACS and analyzed for the EC marker CD31, achieving a high-purity population with 99.11% CD31⁺ cells. Among these, 95,41% co-expressed VE-Cadherin (CD144; VE-Cad-488), confirming endothelial identity.

**Fig. 6:**
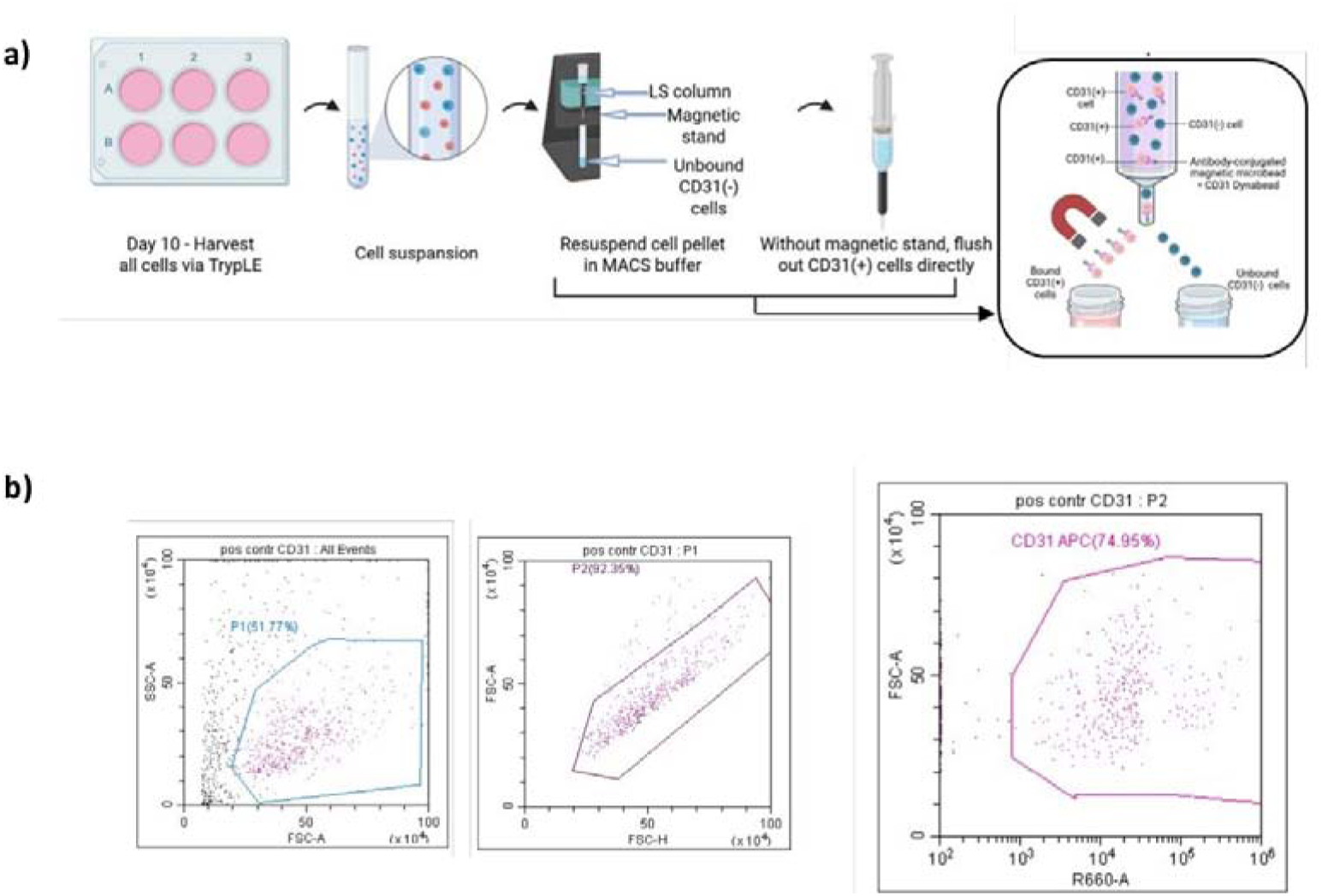
Flow cytometry analysis of endothelial markers following MACS enrichment in APEL media. (a) Magnetic Activation Cell Sorting Dynabeads schematic steps. (b) Cells were cultured in APEL media, sorted by MACS, and after MACS they were analyzed for the EC marker CD31. According to this method, total cell population of 74,95% cells were CD31⁺.

### Morphological characterization of APEL- and BPEL-derived iPSC-EC

Immunofluorescence analysis of iPSC-derived endothelial cells differentiated in APEL medium, characterizing cells, show robust expression of key endothelial markers. Cells were stained for nuclei (DAPI, blue), CD31 (red), and VE-Cadherin (CD144; green). The merged images show co-localization of CD31 and VE-Cadherin at cell–cell junctions, confirming endothelial identity and the formation of characteristic intercellular junctions. These results indicate successful differentiation of functional endothelial cells in APEL medium (Fig. 7). Immunofluorescence analysis of iPSC-derived endothelial cells differentiated in BPEL medium, characterizing cells, show strong expression of EC markers. Cells were stained for nuclei (DAPI, blue), CD31 (red), and VE-Cadherin (CD144; green). The merged images show co-localization of CD31 and VE-Cadherin at cell–cell junctions, confirming EC identity and the formation of proper intercellular junctions. These results demonstrate that BPEL medium efficiently supports the generation of functional ECs comparable to APEL medium (Fig. 7).

**Fig. 7:**
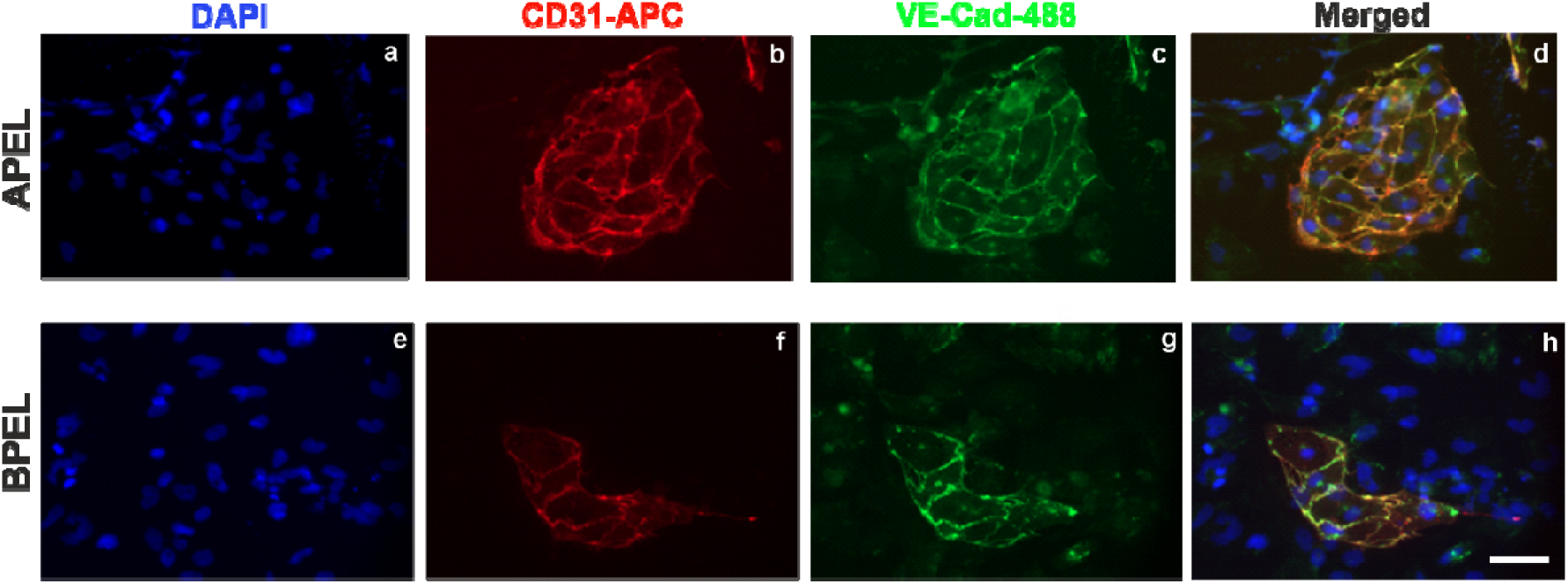
Immunofluorescence analysis of endothelial markers in iPSC-derived ECs in APEL and BPEL. a) EC cultured in APEL media were stained for nuclei (DAPI, blue), b) CD31 (red), and c) VE-Cadherin (CD144; green). The merged image (d) shows co-localization of CD31 and VE-Cadherin at cell–cell junctions, confirming endothelial identity, Day 10. e) EC cultured in BPEL media were stained for nuclei (DAPI, blue), f) CD31 (red), and g) VE-Cadherin (CD144; green). The merged image (h) shows co-localization of CD31 and VE-Cadherin at cell–cell junctions, confirming endothelial identity, Day 10, Scale bar: 50µm

### Functional tub formation of APEL- and BPEL-derived iPSC-ECs

To evaluate the angiogenic functionality of ECs generated using the modified differentiation protocol, iPSC-derived ECs differentiated in either APEL or BPEL medium were subjected to a Matrigel tube formation assay followed by quantitative network analysis using AngioTool (Fig. 8). Representative phase-contrast images demonstrated that ECs generated under both conditions readily formed interconnected capillary-like networks with comparable overall morphology (Fig. 8a,c). Skeletonized AngioTool analysis confirmed the formation of extensive vascular networks in both groups (Fig. 8b,d).

**Fig. 8:**
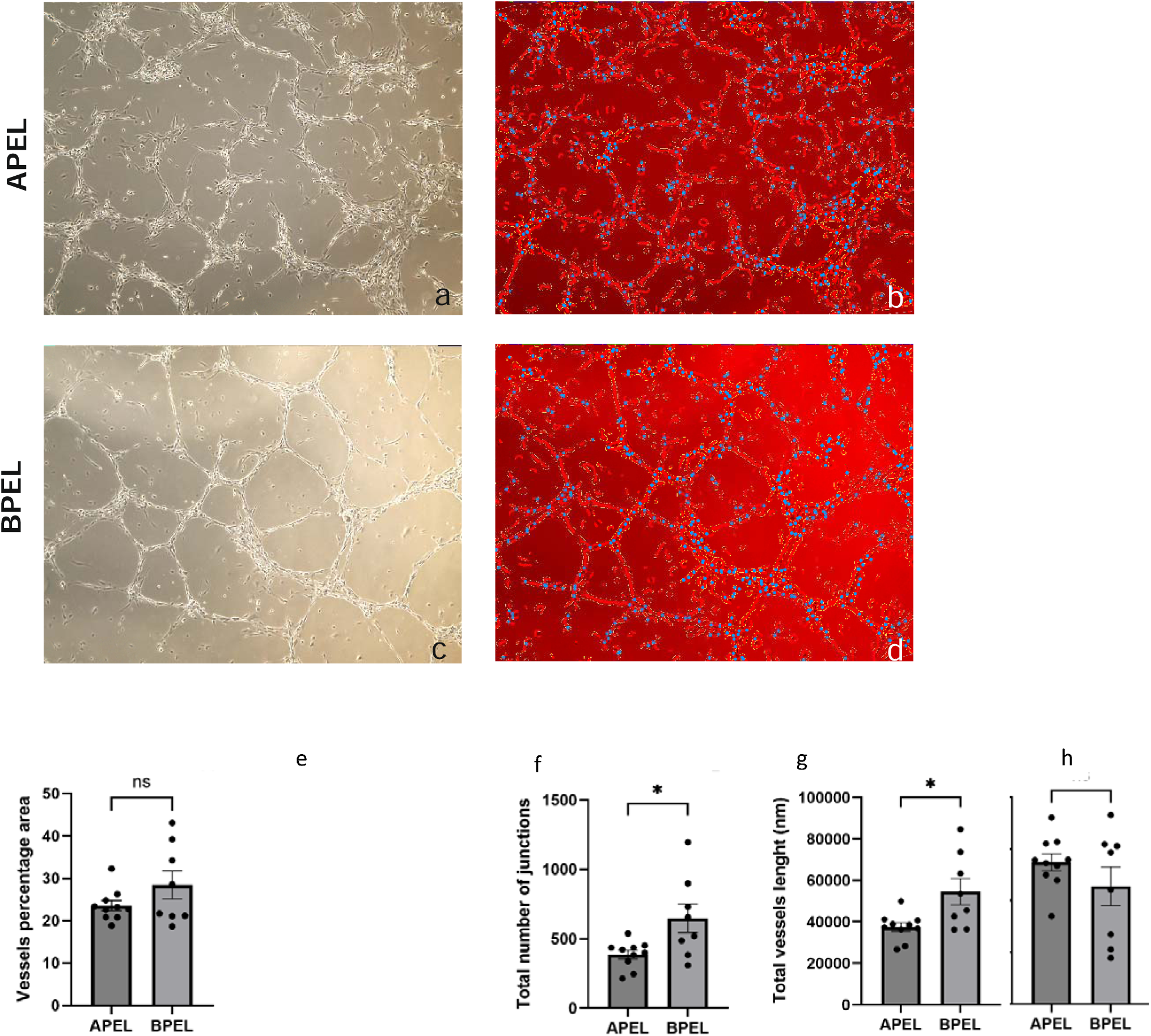
Tube formation analysis via AngioTool. a) hiPSC-derived ECs generating with APEL media, b) AngioTool analysing picture of a, c) hiPSC-derived ECs generating with BPEL media, d) AngioTool analysing picture of c. Phase-contrast micrographs, 4X, day24. Quantitative analysis of angiogenic network parameters including (e) vessel area (%), (f) total number of junctions, (g) total vessel length (mm), and (h) lacunarity. Data are presented as mean ± SEM (n ≥ 3 independent differentiations, each experiment included 2-3 technical replicate wells). Statistical significance was determined using an unpaired Student’s t-test. P < 0.05; ns, not significant.

According to the tube formation analysis of iPSC-derived ECs, total vessel lenght and total number of junctions has increased significantly in BPEL medium-differentiated cells (Total vessel lenght for APEL: 37745 ± 2066 SEM p=0.0318 n=10, for BPEL: 54514 ± 6326 SEM p=0.0318 n=8) (Total number of junctions for APEL: 387.7 ± 30.69 SEM p=0.0409 n=10, for BPEL: 648.5 ± 103.2 SEM p=0.0409 n=8). In addition the vessel percentage area and the lacunarity has also increased in BPEL medium when we compared it with APEL medium-differentiated ECs (Vessel percentage area for APEL: 23.55 ± 1.18 SEM p=0.1987 n=10, for BPEL: 28.43 ± 3.31 SEM p=0.1987 n=8) (Lacunarity for APEL: 0.2742 ± 0.01604 SEM p=0.2841 n=10, for BPEL: 0.2279 ± 0.03756 SEM p=0.2841 n=8) (Fig.8e-f).

### Cost-effectiveness analysis of APEL and BPEL media

To evaluate the economic impact of the differentiation strategy, we first compared the total reagent costs associated with generating hiPSC-derived ECs using APEL and BPEL media (Table 1). The overall reagent cost for a 12-well differentiation was substantially higher for APEL than for BPEL (€686.5 versus €461.5), primarily because of the higher cost of commercially available APEL2 medium. Nevertheless, APEL generated a greater number of CD31⁺ cells following MACS enrichment (approximately 6 × 10⁶ cells) than BPEL (approximately 4 × 10⁶ cells). Consequently, normalization of the total differentiation cost to cell yield revealed a nearly identical cost per 10⁶ differentiated cells (€114.42 for APEL versus €115.37 for BPEL), indicating that the higher initial cost of APEL was largely offset by its increased cell yield (Table 1).

**Table 1:**
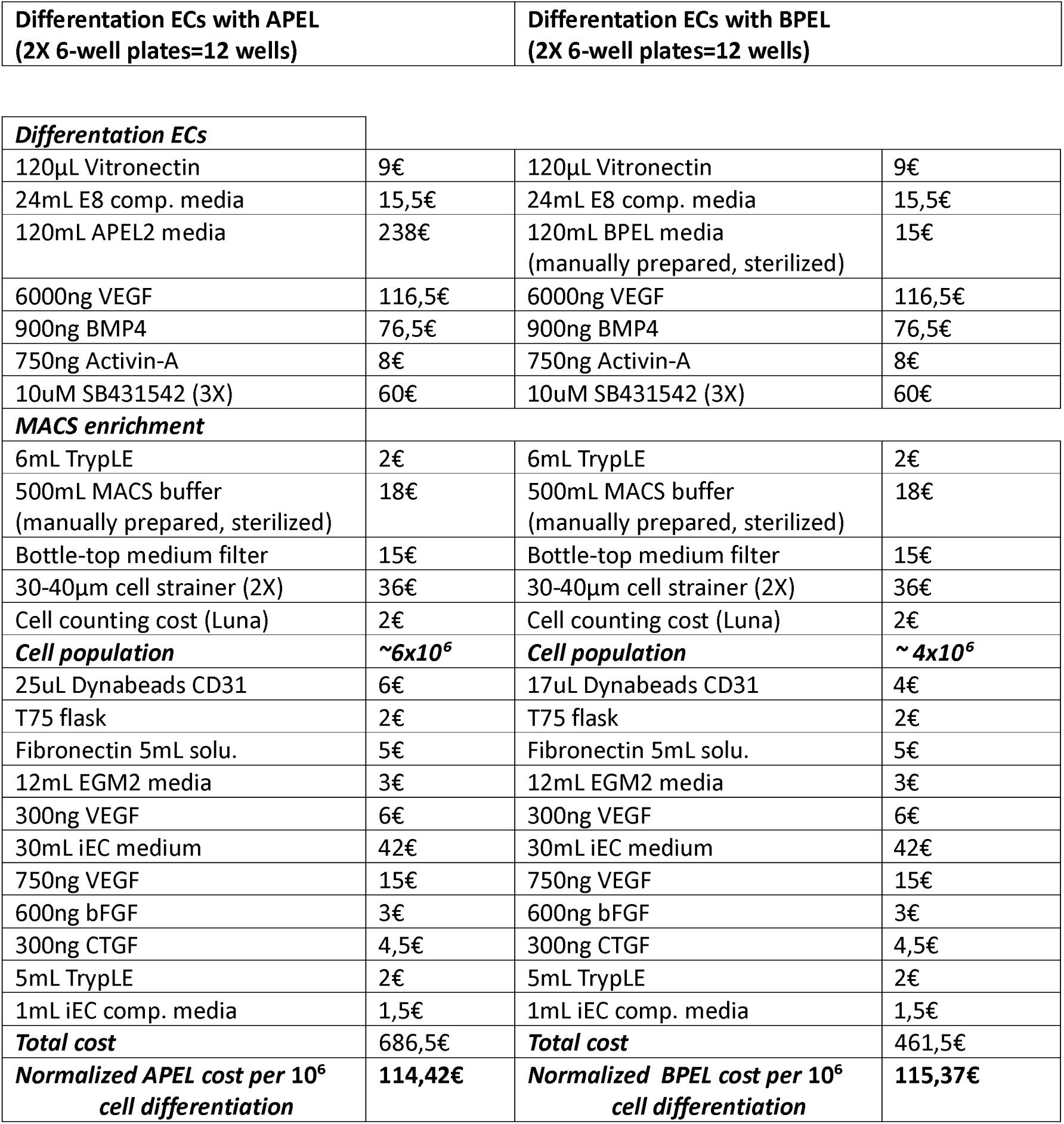
Cost comparison of APEL and BPEL media for the generation of hiPSC-derived ECs. Reagent costs were calculated for differentiation of 12 wells (two 6-well plates) using 2025 prices. The normalized cost per 10⁶ differentiated cells was calculated using the total cell yield obtained at the end of the differentiation protocol.

| Differentiation ECs with APEL<br>(2X 6-well plates=12 wells) | Differentiation ECs with BPEL<br>(2X 6-well plates=12 wells) |
| --- | --- |

**Table 1: Cost comparison of APEL and BPEL media for the generation of hiPSC-derived ECs.**
|  |  |  |  |
| --- | --- | --- | --- |
| **Differentiation ECs** |  |  |  |
| 120µL Vitronectin | 9€ | 120µL Vitronectin | 9€ |
| 24mL E8 comp. media | 15,5€ | 24mL E8 comp. media | 15,5€ |
| 120mL APEL2 media | 238€ | 120mL BPEL media<br>(manually prepared, sterilized) | 15€ |
| 6000ng VEGF | 116,5€ | 6000ng VEGF | 116,5€ |
| 900ng BMP4 | 76,5€ | 900ng BMP4 | 76,5€ |
| 750ng Activin-A | 8€ | 750ng Activin-A | 8€ |
| 10uM SB431542 (3X) | 60€ | 10uM SB431542 (3X) | 60€ |
| **MACS enrichment** |  |  |  |
| 6mL TrypLE | 2€ | 6mL TrypLE | 2€ |
| 500mL MACS buffer<br>(manually prepared, sterilized) | 18€ | 500mL MACS buffer<br>(manually prepared, sterilized) | 18€ |
| Bottle-top medium filter | 15€ | Bottle-top medium filter | 15€ |
| 30-40µm cell strainer (2X) | 36€ | 30-40µm cell strainer (2X) | 36€ |
| Cell counting cost (Luna) | 2€ | Cell counting cost (Luna) | 2€ |
| **Cell population** | **~6x10<sup>6</sup>** | **Cell population** | **~4x10<sup>6</sup>** |
| 25uL Dynabeads CD31 | 6€ | 17uL Dynabeads CD31 | 4€ |
| T75 flask | 2€ | T75 flask | 2€ |
| Fibronectin 5mL solu. | 5€ | Fibronectin 5mL solu. | 5€ |
| 12mL EGM2 media | 3€ | 12mL EGM2 media | 3€ |
| 300ng VEGF | 6€ | 300ng VEGF | 6€ |
| 30mL iEC medium | 42€ | 30mL iEC medium | 42€ |
| 750ng VEGF | 15€ | 750ng VEGF | 15€ |
| 600ng bFGF | 3€ | 600ng bFGF | 3€ |
| 300ng CTGF | 4,5€ | 300ng CTGF | 4,5€ |
| 5mL TrypLE | 2€ | 5mL TrypLE | 2€ |
| 1mL iEC comp. media | 1,5€ | 1mL iEC comp. media | 1,5€ |
| **Total cost** | **686,5€** | **Total cost** | **461,5€** |
| **Normalized APEL cost per 10<sup>6</sup>**<br>**cell differentiation** | **114,42€** | **Normalized BPEL cost per 10<sup>6</sup>**<br>**cell differentiation** | **115,37€** |

Because the ultimate objective of many differentiation protocols is to obtain purified ECs rather than total cell yield, we next compared the cost-efficiency based on the number of CD31⁺/VE-cadherin⁺ (CD144⁺) ECs generated (Table 2). Although APEL produced a higher absolute number of double-positive ECs (4.38 × 10⁶ versus 3.82 × 10⁶), BPEL demonstrated superior cost-efficiency because of its substantially lower reagent cost. The cost per CD31⁺/CD144⁺ cell (€0.0001209 versus €0.0001566), the cost per 10⁶ double-positive cells (€120.93 versus €156.62), and the number of ECs per euro were all more favorable for BPEL (Table 2). These findings suggest that BPEL represents the more economical strategy when the primary objective is the generation of purified ECs.

**Table 2:** Cost-efficiency comparison of APEL and BPEL media for generating double positive CD31⁺/CD144⁺ hiPSC-derived ECs. Costs were normalized to the number of CD31⁺/CD144⁺ cells obtained after enrichment.

| Metric | APEL | BPEL | Interpretation |
| --- | --- | --- | --- |
| Total cost | €686,5 | €461,5 | Lower price in BPEL |
| CD31<sup>+</sup>/CD144<sup>+</sup> number | 4.383.000 | 3.816.400 | Higher cells in APEL |
| Cost per CD31<sup>+</sup>/CD144<sup>+</sup> cell | €0.0001566 | €0.0001209 | Lower price in BPEL |
| Cost per 10<sup>6</sup> CD31<sup>+</sup>/CD144<sup>+</sup> cells | €156,62 | €120,93 | Lower price in BPEL |
| CD31<sup>+</sup>/CD144<sup>+</sup> cells per €1 | 6 | 8 | Higher cells in BPEL |

Finally, we evaluated cost-efficiency using the total number of CD31⁺ cells recovered after MACS enrichment (Table 3). Although APEL required a higher initial investment, its greater overall cell yield resulted in a slightly lower cost per CD31⁺ cell (€0.0001144 versus €0.0001154) and a marginally lower cost per 10⁶ CD31⁺ cells (€114.42 versus €115.38). Likewise, the number of CD31⁺ cells generated per euro was slightly higher with APEL. These findings indicate that APEL may be the preferred option when maximizing total EC production is the primary objective, whereas BPEL offers superior cost-efficiency for generating highly endothelial-specific (CD31⁺/CD144⁺) cell populations (Table 3).

**Table 3:** Cost-efficiency comparison of APEL and BPEL media based on total cell yield after enrichment. Costs were normalized to the total number of cells obtained after enrichment, based analyse both CD31⁺ and CD31⁻ cell populations.

| Metric | APEL | BPEL | Interpretation |
| --- | --- | --- | --- |
| Total cost | €686,5 | €461,5 | Lower price in BPEL |
| Total CD31<sup>+</sup>cells | 6.000.000 | 4.000.000 | Higher cells in APEL |
| Cost per CD31<sup>+</sup>cell | €0.0001144 | €0.0001154 | Lower price in APEL |
| Cost per 10<sup>6</sup> CD31<sup>+</sup>cells | €114,42 | €115,38 | Lower price in APEL |
| CD31<sup>+</sup> Cells per €1 | 8 | 8 | Higher cells in APEL |

## Supplement

**Table S1:** Reagents and Materials.

| Material / Reagent | Company | Catalog Number |
| --- | --- | --- |
| Vitronectin | Gibco | 15134499 |
| Essential 8 (E8) Medium | Gibco | A2858501 |
| ROCK inhibitor (Y-27632) | Tocris | 1254/1 |
| APEL2 Medium | STEMCELL Technologies | 0-5270 |
| VEGF165 | PeproTech | 100-20-50UG |
| BMP4 | Sigma-Aldrich | 130111168 |
| Activin A | R&D Systems | 338-AC-010 |
| CHIR99021 | R&D Systems | 4423/10 |
| 1-Thioglycerol (a-MTG) | Sigma Aldrich | M6145-25ml |
| SB431542 | STEMCELL Technologies | 72232 |
| BPEL Medium components: |  |  |
| IMDM (no phenol red) | Gibco | 21056023 |
| Ham's F12 + GlutaMAX | Gibco | 31765027 |
| Protein-Free Hybridoma Medium | Gibco | 12040077 |
| Chemically Defined<br>LipidConcentrate (CD<br>Lipids) | Gibco | 11905031 |
| ITS-X Supplement (100X) | Gibco | A4000046501 |
| GlutaMAX | Gibco | 35-050-061 |
| Primocin | InvivoGen | ant-pm-05 |
| BSA | Sigma-Aldrich | A3311 |
| Polyvinyl alcohol (PVA) | Sigma-Aldrich | P8136-250G |
| L-Ascorbic Acid | Sigma-Aldrich | A8960-5G |
| TrypLE Express | Gibco | 12604013 |
| CD31 Dynabeads | Invitrogen | 11155D |
| MACS Buffer Components (BSA, EDTA, PBS) | Various (Sigma Aldrich, Thermo Fisher, Sigma Aldrich) | A3311,15575020, P4417-100TAB |
| Cell Strainer (35-40µm) | Falcon / Corning | 352340 |
| Fibronectin | Sigma-Aldrich | F2006 |
| EGM-2 BulletKit | PromoCell | C-22011 |
| Revitacell Supplement | Gibco | A2644501 |
| VascuLife Medium Kit | Lifeline Cell Technology | LL-0005 |
| bFGF | StemCell Technologies | 78003 |
| CTGF | Gibco | 120-19-5UG |
| Primary Antibodies: |  |  |
| CD31-APC | Miltenyi Biotech | 13011067 |
| VE-Cad-488 (CD144) | Invitrogen | 53144941 |

## Author contributions

PNK, and LK. performed research and analyzed data, ZH designed and supervised research, and ZH, PNK, and LK wrote the manuscript.

## Ethics statements

Not applicable.

## Acknowledgements

We thank Sarah de Jong and Yoanna Petrova for their scientific contributions. Schematic figures were created by biorender.com.

